# Environmental Fungi Modulate the Vaginal Mycobiome and Cervical Disease Progression in Hispanic Women

**DOI:** 10.64898/2026.01.21.700834

**Authors:** Filipa Godoy-Vitorino, Daniela Vargas-Robles, Benjamin Bolaños-Rosero, Natalia Pagán-Zayas, Andrea Cortés-Nazario, Kara Wiggin, Sarah Allard, Josefina Romaguera, Jack A. Gilbert

## Abstract

The vaginal mycobiome, though a minor component of the cervicovaginal ecosystem, plays a crucial role in reproductive health and disease. However, its composition and interactions with bacterial communities remain poorly understood, particularly among Hispanic women, who experience disproportionately high rates of Human papillomavirus (HPV) infection and cervical cancer. We characterized the vaginal mycobiota across reproductive stages and examined its associations with cervical disease, HPV status, and bacterial community state types (CSTs) in 86 Hispanic participants from Puerto Rico using ITS1 amplicon sequencing. Amplicon sequence variants were inferred with QIIME2/DADA2 and taxonomically classified using the UNITE database, with diversity and discriminant taxa analyses applied to explore clinical and microbial associations. We detected 173 fungal Species Hypotheses, dominated by *Candida albicans*, *Agaricomycetes* sp., *Scopuloides dimorpha*, and *Hortaea werneckii*. While fungal composition did not differ significantly by reproductive stage, non-pregnant individuals exhibited greater inter-individual variability. Alpha diversity was reduced in high-grade squamous intraepithelial lesions compared with low-grade or normal cytology, and *Candida parapsilosis* prevalence was elevated in low-grade lesions. CST III, characterized by *Lactobacillus iners* dominance, showed greater dispersion variance than other CSTs. Collectively, these findings reveal a diverse vaginal mycobiome with stage- and disease-specific features, and a notable contribution of environmental fungi that may influence cervical pathogenesis. This work provides foundational insight into cervicovaginal fungal ecology in a high-risk Hispanic population and highlights the importance of integrating bacteriome–mycobiome analyses in women’s health research.

## Introduction

The vaginal microbiome is a complex and dynamic ecosystem that plays a critical role in maintaining female reproductive health [1, 2]. While bacterial communities have been extensively studied, the fungal component—known as the vaginal mycobiome—remains relatively unexplored [3, 4]. Early research suggested that *Candida albicans* is the dominant fungal species in the vagina, yet recent advances in sequencing technologies have revealed a much broader fungal diversity, with potential implications for gynecological health and disease [5]. Fungi, despite being a minor component of the human microbiome, have been found in several body sites, including the skin [6], gut [7], lung [8], oral cavity [9, 10], nose [11] and reproductive tract [3, 4], where they interact with bacteria, the host immune system, and environmental factors to influence health and disease. In the vaginal environment, fungi are thought to be present in low abundance under normal conditions, but they can become opportunistic pathogens under specific circumstances, such as immune dysregulation or microbial imbalance [5, 12].

In the cervicovaginal niche, the mycobiome has long been overshadowed by the study of bacterial communities, mainly due to early reliance on culture-based methods that underestimated fungal diversity [5]. However, next-generation sequencing and bioinformatics approaches have increasingly demonstrated that the vaginal mycobiota consists of more than just *Candida* species. Other fungi, including *Saccharomyces cerevisiae*, *Malassezia restricta*, and various filamentous fungi, have been detected, though their roles remain poorly understood [3, 4]. These fungi may exist in a commensal state, contributing to vaginal homeostasis, or may play a more pathogenic role in conditions such as vulvovaginal candidiasis [13], bacterial vaginosis (BV) [14], and sexually transmitted infections [15]. The interplay between bacteria and fungi in the cervix is an emerging area of research, particularly in the context of HPV infections. *Candida albicans*, the most well-characterized vaginal fungal species, has been shown to enhance inflammation and disrupt epithelial integrity, potentially facilitating HPV persistence and cervical lesion progression [12]. Moreover, studies have demonstrated that bacterial dysbiosis, particularly a decrease in *Lactobacillus* species and an overgrowth of anaerobes such as *Gardnerella vaginalis*, is often accompanied by shifts in fungal composition. The presence of fungal species, such as *Malassezia* and *Saccharomyces*, in HPV-positive cervical samples raises questions about how fungi interact with bacteria and the host immune system to influence HPV persistence and disease progression [5, 16]. Additionally, fungal colonization has been linked to increased levels of pro-inflammatory cytokines such as interleukin-17A (IL-17A), which may contribute to a more permissive environment for viral persistence and neoplastic transformation [12, 17]. Given the high incidence of HPV and cervical cancer among Puerto Rican Hispanics [18] understanding the mycobiome in this population is crucial for advancing individuals’ health research.

Hispanic women living in Puerto Rico have non-optimal bacterial communities distinct from other populations [19]. Research indicates that the cervicovaginal microbiota of Hispanics in Puerto Rico is commonly dominated by *Lactobacillus iners* and anaerobic bacteria rather than by the protective *Lactobacillus crispatus*. These typical Puerto Rican profiles, often enriched with BV-associated bacteria such as *Atopobium vaginae* and *Gardnerella vaginalis*, has been linked to cervical intraepithelial neoplasia (CIN), particularly CIN3 precancerous lesions [19, 20]. Interestingly, while no specific bacterial biomarkers were identified for HPV infections, fungal diversity was elevated in cases of high-risk HPV, suggesting a potential role for fungi in modifying disease susceptibility and immune responses [4].

In this study, we analyze the vaginal mycobiota of 86 Hispanic individuals from Puerto Rico across different reproductive stages (pregnant, non-pregnant, and menopausal) and examine mycobiome diversity associations with cervical disease and HPV status among non-pregnant individuals. We identify key taxa associated with cervical lesions and explore their potential interactions with bacterial community state types (CSTs). By addressing this knowledge gap, we hope to contribute to a more comprehensive understanding of the cervicovaginal microbiome and its role in individuals’ reproductive health, particularly in these high-risk populations.

## Methods

### Study Design and Participant Sample Collection

This cross-sectional study included adult women attending gynecology and colposcopy appointments at the University of Puerto Rico (UPR) and San Juan City clinics, located in the metropolitan area of San Juan, Puerto Rico. Participants who met the inclusion criteria and did not fulfill any exclusion criteria were invited to participate. Exclusion criteria were: (1) a history of regular urinary incontinence, (2) prior treatment for or suspicion of toxic shock syndrome, (3) current candidiasis, (4) active urinary tract infection, (5) active sexually transmitted infection (STI), and (6) cervicovaginal irritation at the time of screening. Only asymptomatic participants were enrolled, and STI testing was not routinely performed in this population. Women who had received antibiotic treatment in at least two months prior to enrollment were not excluded, as they represented a small proportion of the study cohort and were evenly distributed across groups.

Data and biological samples were collected between 2018 and 2021 from participants aged 21 to 60 years. The study protocol was reviewed and approved by the Institutional Review Board (IRB) of the UPR Medical Sciences Campus (protocol #1050114, June 2014) and the Ethics Committee of San Juan City Hospital, with biosafety approval under protocol #94620. All participants provided written informed consent and signed HIPAA authorization forms after receiving verbal and written explanations of study procedures, risks, and benefits as per the Declaration of Helsinki. All study personnel were certified in research ethics and compliance training, including CITI RCR, Social and Behavioral Research Best Practices, HIPAA, and NIH Protection of Human Subjects. Participants completed a detailed metadata questionnaire capturing demographic data (age, birthplace, employment, education), sexual history (age at sexual debut, number of partners), health and medical history, antibiotic and vitamin use, and body mass index (BMI).

A sterile speculum was used to visualize the cervix. Cervical samples were obtained by gently gentle rotations in a circular motion in the posterior fornix - a sampling site selected to optimize HPV detection and provide a representative profile of the cervicovaginal microbiota as in previous studies [19]-using sterile Catch-All™ Specimen Collection Swabs (Epicentre Biotechnologies, Madison, WI) and immediately placed in MoBio bead tubes containing MoBio PowerSoil™ buffer (MoBio, Carlsbad, CA). Swabs were agitated for approximately 20 seconds in 750 μL of buffer before removal. Additionally, ∼10 mL of cervical lavage fluid was collected by flushing the vaginal canal with Pure certified nuclease-free water (Growcells, Irvine, CA, USA) for future analyses. Lavage samples were transferred to 15 mL tubes, and pH was measured using Hydrion pH paper strips. All samples were labeled, placed on ice for up to 4 hours, transported to the laboratory, and stored at -80 °C until nucleic acid extraction and subsequent molecular analyses were performed. All processing and PCR procedures were performed in a single laboratory (FGV) to minimize technical variation.

### DNA Extraction and 16S rRNA Gene Sequencing and metadata

Genomic DNA was extracted from posterior fornix cervical swabs using the Qiagen PowerSoil Kit (QIAGEN LLC, Germantown Road, Maryland, USA), following an optimized protocol described previously [16]. No human DNA depletion or microbial/viral DNA enrichment steps were performed. Briefly, PowerBead tubes were homogenized for 10 minutes at 3,200 rpm using a Vortex-Genie 2 G560 (Scientific Industries, Inc., NY). Solutions C2 and C3 (100 μL each) were combined and vortexed for 5 seconds to facilitate cell lysis. DNA was eluted in 100 μL of sterile PCR-grade water pre-warmed to 55 °C, which was allowed to remain on the filter for 5 minutes before the final centrifugation step to maximize yield. DNA concentrations were quantified using the Qubit® dsDNA High Sensitivity Assay (Thermo Fisher Scientific, Waltham, MA), with yields ranging from 5–100 ng/μL. Genomic DNA was shipped to an external sequencing facility with average concentrations of 10–30 ng/μL.

For 16S rRNA sequencing, extracted DNA was normalized to 4 nM during library preparation. The hypervariable V4 region of the 16S rRNA gene (∼291 bp) was amplified using universal bacterial primers 515F (5’-GTGCCAGCMGCCGCGGTAA-3’) and 806R (5’-GGACTACHVGGGTWTCTAAT-3’), as established by the Earth Microbiome Project (http://www.earthmicrobiota.org/emp-standard-protocols/16s/) [21]. Sequencing was outsourced to Argonne National Laboratory (Illinois, USA) and performed on an Illumina MiSeq platform using a 2×250 bp paired-end kit, and bacterial diversity data was previously reported [19]. We only used CST groups as determined by VALENCIA [22].

Clinical data, including cervical cytology and pathology results, were obtained from patient medical records. Cytological findings were classified according to the Bethesda system as: negative for intraepithelial lesion or malignancy (NILM), low-grade squamous intraepithelial lesion (LGSIL), or high-grade squamous intraepithelial lesion (HGSIL) [23]. Due to a low number of samples, patients with a cytology result of atypical squamous cells of undetermined significance (ASC-US) categorized as either HPV negative or HPV-lr positive exclusively (n = 4) were reclassified as NILM. Patients identified as ASC-US with HPV positive (n = 2) were classified as LGSIL.

All sequencing reads and associated metadata were uploaded to QIITA [24] (study ID: 12871, https://qiita.ucsd.edu/study/description/12871). Raw ITS sequencing data were deposited in the European Nucleotide Archive (ENA) under accession number ERP136546.

### HPV Genotyping and Cytologic Evaluation

HPV genotyping was performed using SPF10-LiPA, a highly sensitive short polymerase chain reaction-fragment assay (Labo Biomedical Products, Rijswijk, The Netherlands), based on Innogenetics-licensed technology. Amplified products were subjected to reverse hybridization for genotype identification by comparison to standardized kit controls as previously described [19]. This assay enables detection of 25 mucosal HPV genotypes including eleven low-risk (HPV-lr): 6, 11, 34, 40, 42, 43, 44, 53, 54, 70, and 74; and fourteen high risk (HPV-hr): 16, 18, 31, 33, 35, 39, 45, 51, 52, 56, 58, 59, 66, 68/73. Genotyping results were categorized into three groups: (1) HPV-hr only, (2) HPV-lr only, and (3) HPV-negative (**Table 1**).

**Table 1.** Demographic and clinical characteristics of the studied population.

| Variable | level | Pregnant | Non-pregnant | Menopause | p value & | Overall |
| --- | --- | --- | --- | --- | --- | --- |
| n |  | 9 | 63 | 14 |  | 86 |
| Age (mean (SD)) |  | 28.00 (4.92) | 36.17 (8.84) | 54.00 (5.19) | <0.001 * | 38.22 (10.89) |
| BMI (mean (SD)) |  | 32.21 (10.55) | 28.85 (7.53) | 26.67 (4.42) | 0.231 | 28.85 (7.53) |
| Bacterial CST by 16S (%) | I | 0 ( 0.0) | 11 (17.5) | 3 ( 21.4) | 0.016 * | 14 (16.3) |
|  | II | 0 ( 0.0) | 3 (4.8) | 1 ( 7.1) |  | 4 (4.7) |
|  | III | 6 (66.7) | 19 (30.2) | 3 ( 21.4) |  | 28 (32.6) |
|  | IV | 1 ( 11.1) | 29 (46.0) | 7 ( 50.0) |  | 37 (43.0) |
|  | V | 2 (22.2) | 1 ( 1.6) | 0 ( 0.0) |  | 3 (3.5) |
| Cervical Disease Status (%) | Negative | 1 ( 11.1) | 43 (68.3) | 14 (100.0) | <0.001 * | 58 (67.4) |
|  | Positive | 8 (88.9) | 20 (31.7) | 0 ( 0.0) |  | 28 (32.6) |
| Cervical Disease Type (%) | NILM | 1 ( 11.1) | 43 (68.3) | 14 (100.0) | 0.001 * | 58 (67.4) |
|  | LGSIL | 4 (44.4) | 10 (15.9) | 0 ( 0.0) |  | 14 (16.3) |
|  | HGSIL | 4 (44.4) | 10 (15.9) | 0 ( 0.0) |  | 14 (16.3) |
| HPV Status (%) | Negative | 2 ( 22.2) | 24 (38.1) | 7 ( 50.0) | 0.408 | 33 (38.4) |
|  | Positive | 7 (77.8) | 39 (61.9) | 7 ( 50.0) |  | 53 (61.6) |
| HPV types (%) | Negative | 2 ( 22.2) | 24 (38.1) | 7 ( 50.0) | 0.577 | 33 (38.4) |
|  | HPV-lr exclusively | 0 ( 0.0) | 3 ( 4.8) | 1 ( 7.1) |  | 4 (4.7) |
|  | HPV-hr | 7 (77.8) | 36 (57.1) | 6 (42.9) |  | 49 (57.0) |
| HPV type richness total (mean (SD)) |  | 2.00 (0.82) | 1.92 (1.06) | 2.57 (1.40) | 0.352 | 2.02 (1.08) |
| HPV-hr type richness (mean (SD)) |  | 1.71 (0.76) | 1.47 (0.61) | 2.33 (1.51) | 0.049 * | 1.61 (0.81) |
| HPV-lr type richness (mean (SD)) |  | 1.00 (0.00) | 1.29 (0.47) | 1.33 (0.58) | 0.689 | 1.27 (0.46) |
| Antibiotic use last 2 months (%) | No | 9 (100.0) | 60 (95.2) | 14 (100.0) | 0.567 | 83 (96.5) |
|  | Yes | 0 ( 0.0) | 3 ( 4.8) | 0 ( 0.0) |  | 3 (3.5) |

& p value is calculated using Fisher Exact Test and Kruskal Wallis/T test for categorical and numerical variables respectively

### Sequencing the ITS1 genes

Fungal community profiling was outsourced at the Argonne National Lab and performed by amplifying the internal transcribed spacer (ITS) region using a dual-indexed paired-end sequencing approach on the Illumina platform. The EMP.ITSkabir primer set, designed for broad coverage of fungal and other microbial eukaryotic lineages, was used to target the ITS1 region [25, 26]. The forward primer (ITS1f) is located 38 bp upstream of the ITS1 region [26] and was synthesized with a 5′ Illumina adapter and linker sequence (AATGATACGGCGACCACCGAGATCTACAC GG CTTGGTCATTTAGAGGAAGTAA). The reverse primer (ITS2), identical to that described by White et al. (1990) [26], included the reverse complement of the 3′ Illumina adapter, a unique 12-bp Golay barcode for sample multiplexing, and a linker sequence (CAAGCAGAAGACGGCATACGAGATNNNNNNNNNNCGGCTGCGTTCTTCATCGATGC). PCR reactions were performed under standard cycling conditions optimized for fungal ITS amplification, and resulting amplicons were pooled, purified, and sequenced using paired-end chemistry. This protocol has been shown to enhance detection sensitivity and taxonomic resolution of fungal communities while minimizing amplification bias [27].

### Sequence processing

Raw data were processed using QIIME2 (version 2023.7). Amplicon sequence variants (ASVs) were inferred using DADA2. Taxonomic classification was conducted using the UNITE dynamic database (version 9, 25.07.2023) and a pre-trained Naïve Bayes classifier in QIIME2. The UNITE database assigns each fungal ASV to a Species Hypothesis (SH), which represents a cluster of sequences with ≥97% similarity and is used as a proxy for species-level identification in fungal ecology. Each SH is associated with a unique identifier (e.g., “SHXXXXXXX”) and serves as a standardized taxonomic unit when formal species names are unavailable or uncertain. These SH identifiers were used as the species-level taxonomic resolution for all downstream statistical analyses.

The ASV table was then imported into R (version 4.1.2) for downstream filtering. In R, unclassified ASVs at Phylum, Class and Order levels we removed. Additionally, sequences corresponding to chloroplasts, mitochondria, archaea, and any non-Fungi sequence (k__Eukaryota_kgd_Incertae_sedis, k__Protista) were removed. A subset of the unclassified ASVs with the highest read abundance was manually curated using NCBI BLAST to confirm taxonomic assignments. We removed samples with fewer than 100 reads, taxa with fewer than 2 sequences or with no SH classification. No rarefaction was applied due to the low number of reads for an important number of samples, that would cause to discard a considerable number of samples.

### Statistical Analyses

The *phyloseq* package [28] in R was used to conduct vaginal mycobiota diversity analyses. The models assessed reproductive stage (Pregnant, Non-Pregnant, and Menopausal) across all patients, while cervical disease status (HGSIL, LGSIL, and NILM), HPV type status (HPV-hr and Negative), and CST from bacterial microbiome (CST-I, CST-III, and CST-IV), only among Non-Pregnant individuals. Age and BMI were included as confounding variables in all models (alpha/beta diversity and discriminant taxa analysis), except for the reproductive stage variable, which did not include Age due to high collinearity with the reproductive stage variable.

Mycobial alpha and beta diversity were estimated at the SH level. To measure alpha diversity Shannon index was calculated and set as the dependent variable. Independent variables (Group or Cervical disease or HPV status, or CST) were evaluated using linear regression models. Normality in the distribution of the Shannon metric was verified and confirmed using “shapiro.test” R function. Shannon’s normality made it suitable to run linear regression models without any transformation. The "step" function in R was used to determine the best-fitted model by systematically eliminating non-relevant variables. We used “emmeans” and the “pairs” R function to conduct pairwise analysis of specific variables.

Beta diversity was analyzed by computing Bray-Curtis and Jaccard distances, followed by Permutational Multivariate Analysis of Variance (PERMANOVA[29]) with the “adonis2” [30] R function for community composition differences and Permutational Analysis of Multivariate Dispersions (PERMDISP [31]) to evaluate inter-individual dispersion with “betadisper” [30] R function. Non-metric Multidimensional Scaling (NMDS) was employed to ordinate beta diversity. Specifically, for the Reproductive Stage variable, two outliers (SampleID: 2031, 2113b) were excluded to enhance sample dispersion visualization.

Discriminant taxa analyses were conducted at the Species and Genus levels. Discriminant taxa to each study variable were assessed using MaAsLin2’s [32], including only the taxa present in at least 10% of the samples and with a relative abundance of at least 10%. Prevalences analyses were performed at the Species level using Fisher Exact Test. P values were adjusted for multiple comparisons using the false discovery rate method in all analyses that involved generating multiple p values (e.g., pairwise comparisons, multiple prevalence calculations, or discriminant taxa analyses). All analyses files and scripts are available at github https://github.com/UPR-MicrobiomeLab/ITS_86_HPV.

## Results

### Study population

The cohort included 86 Hispanic individuals: Menopausal (n = 14), Non-Pregnant (n = 63), and Pregnant (n = 9). Age significantly differed across groups (p < 0.001), while BMI did not (p = 0.231). Bacterial CSTs prevalence varied by group (p = 0.016), with Pregnant individuals mainly exhibiting CST-III, and the other groups dominated by CST-IV. Pregnant individuals showed the highest prevalence of cervical disease (88.9%, p < 0.001), specifically of high-grade lesions (HGSIL, 44.4%, p = 0.001). HPV infection rates were similar among groups (p = 0.408), and most of them were high-risk HPV (HPV-hr, 92.5%). HPV-hr type richness (count of different HPV types in a sample) was significantly greater in Menopausal individuals (p = 0.049). Antibiotic use in the last two months was infrequent and similar across groups (p = 0.567) (**Table 1**).

### Fungal taxonomic diversity

Among 102 sequenced samples, 16 were removed: 15 had fewer than 100 sequences, and one had a single ASV classified as Protista (SampleID: 2302). We finally analyzed 86 samples/individuals. Across these, we found 457 ASVs, corresponding to 173 Species Hypothesis (SH), from which 120 (69%) occurred exclusively in single samples and were not shared between samples, indicating a highly heterogeneous and individualized fungal community with few recurrent taxa. The remaining 31% were shared by two or more samples **(Supp text S1)**.

The 173 SHs represented 125 distinct genera, but seven samples (SampleID: 2218, 2031, 2113b, 2278, 2302, 2306, 2308) showed an absolute dominance of a single SH, reflecting extremely low-complexity communities. Across the entire dataset, only five SHs were detected in 10 or more samples: the human-associated yeast *Candida albicans* and the environmental yet human-associated yeast *Hortaea werneckii* (both Ascomycota), as well as three environmental filamentous fungi from the class Agaricomycetes - *Agaricomycetes* sp., *Scopuloides dimorpha*, and *Trametes cubensis* (**Supp text S1)**.

Two individuals who presented only unclassified SHs and were removed then the analyses were performed at this specific taxonomic level (SampleID: 2128b, with the absolute dominance of one ASV with classification available up to Genus: Polyporales_sp and, SampleID: 2292, with composed only by two ASVs classified as g Aspergillus or f Teratosphaeriaceae).

*Candida albicans* was present in 45.3% of the samples (39 out of 86) and had the highest mean relative abundance of 43.1%, when present. The next most abundant SH was the non-human associated *Agaricomycetes_sp* with an overall prevalence of 50.0% (43 out of 86) and a mean relative abundance of 29.8%, when present (**Fig. 1**). We identified a total of 11 human-associated taxa and 162 non-human-associated SHs (**Fig. 1**).

**Figure 1.**
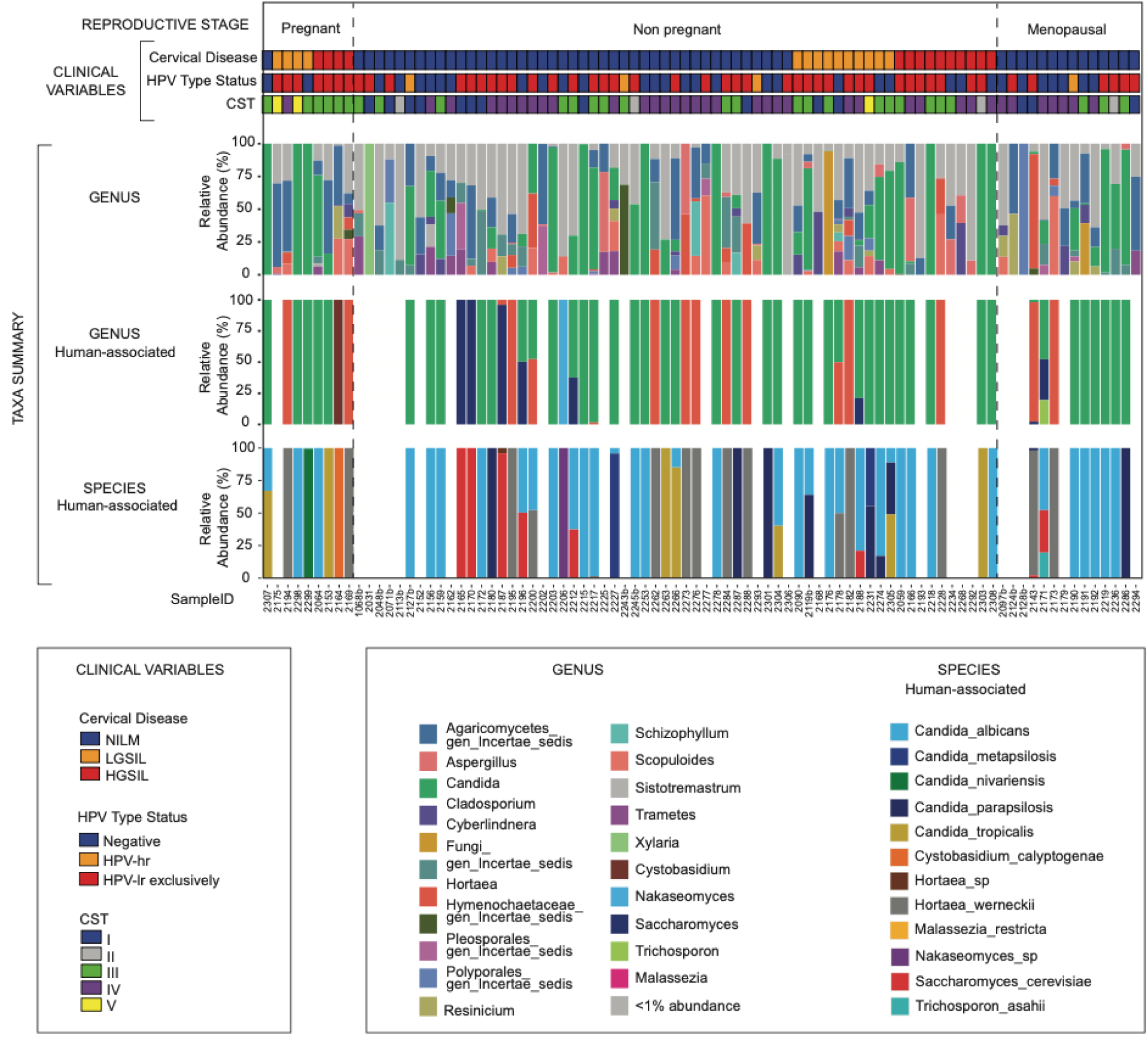
Taxonomic composition and clinical metadata of vaginal samples across reproductive stages. Each column represents one sample, grouped into Pregnant, Non-Pregnant, and Menopausal stages (separated by dashed vertical lines). The top panel shows clinical metadata including cervical disease status (NILM, LGSIL, HGSIL), HPV type status (Negative, HPV-hr, HPV-lr exclusively), and Community State Types (CST I–V). The taxonomic summary displays the relative abundance (%) of fungal taxa at three levels: all detected genera (GENUS), human-associated genera (GENUS Human-associated), and human-associated species (SPECIES Human-associated). Taxa with low relative abundance are grouped as "<1% abundance". Color assignments are consistent across panels and correspond to the legends below. Sample IDs are displayed on the x-axis.

### Reproductive stage groups analysis (Pregnant, Non-Pregnant, and Menopausal comparison)

We compared vaginal fungal diversity and composition among reproductive stages (Pregnant, Non-Pregnant, and Menopausal) including all individuals and then cervical disease (LGSIL, HGSIL, and NILM), HPV type status (HPV-hr, Negative) and vaginal 16S CST (CST I, III, IV), including only Non-Pregnant individuals. Reproductive stages were significantly associated with differences in beta dispersion (Jaccard, p = 0.004; Bray-Curtis, p = 0.046). Specifically, Non-Pregnant women showed a significantly greater inter-individual compositional heterogeneity than Pregnant women (p adj = 0.025, Jaccard, **Fig 1B**), while Menopausal individuals showed a marginally trend toward greater heterogeneity compared to Pregnant women (p adj = 0.074, Jaccard, PERMDISP test, **Fig. 1B**). The reproductive stages did not have a significant difference in fungal composition (PERMANOVA, p > 0.723, **Fig 1A, Supp text S2**). No significant differences in alpha diversity were observed across reproductive stages (p > 0.050, Shannon index, linear model, **Supp text S3**). Discriminant taxa and taxa prevalence analyses suggested that no taxa were significantly associated (p < 0.050, p adj < 0.05) with the reproductive state groups after p value adjustment for multiple testing. However, before p value adjustment, the relative abundance of the genus *Recinicium* (11/86) was greater in Menopausal individuals (p = 0.027, p adj = 0.562; MaAsLin2, **Fig 2C, Supp text S4**) and *Candida tropicalis* prevalence appeared marginally higher in Pregnant (33.3%) than in Menopausal (0%) or Non-Pregnant individuals (7.8%) (*p* = 0.051, p adj = 0.568, Fisher’s Exact Test; **Supp text S5**), though these associations did not remain significant after adjustment.

**Figure 2.**
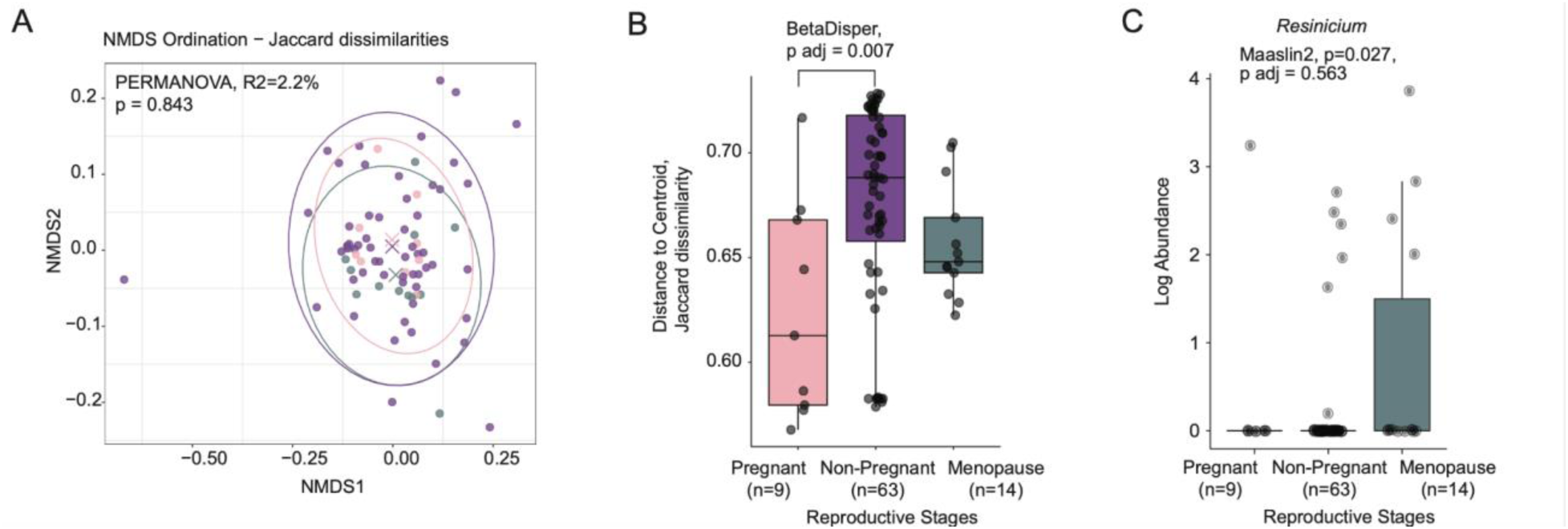
Vaginal mycobiome diversity by Reproductive Stage. **A.** Ordination indicating centroids with an X symbol and 95% confidence interval ellipse by using Jaccard dissimilarity, analyzed by PERMANOVA. **B.** Distance to centroid using Jaccard dissimilarity, analyzed using PERMDISP test and its pairwise form. **C.** *Resinicium* genus relative abundance with logarithmic transformation, identified by MaAsLin2 as significantly associated taxa prior to p value adjustment for multiple comparisons.

### Cervical disease and HPV status among Non-Pregnant individuals

Cervical disease was evaluated among Non-Pregnant individuals due to the limited sample size and substantial physiological differences between the other groups (Pregnant and Menopausal, **Table 1**). While beta diversity and dispersion showed no significant difference between cervical disease groups (LGSIL, HGSIL, and NILM, p > 0.356, Bray-Curtis and Jaccard, **Supp text S2**), alpha diversity was significantly lower in HGSIL compared to LGSIL (p = 0.037) or NILM (p = 0.037), and LGSIL had greater alpha diversity than NILM (p=0.049, Estimated Marginal Means, **Fig. 1C, Supp text S3**). *Candida parapsilosis* (7/63) was at a greater proportion in LGSIL than NILM, but not significantly after p value adjustment (p = 0.011, p adj = 0.497, MaAsLin2). Although no taxa remained significant after correction for multiple comparisons, prior to p adjustment *Candida parapsilosis* was more prevalent in LGSIL samples (40.0%, 4 of 10) compared with NILM (11.1%, 7 of 43; p = 0.020) and HGSIL (0%, 0 of 10; p = 0.038; Fisher’s Exact Test; **Supp text S6**).

HPV status did not show any significant difference for either beta using PERMANOVA or PERMDISP or alpha diversity (HPV-hr vs HPV negative, p > 0.327, **Supp text S2, S3**). The category “exclusively HPV-lr” was removed from this comparison due to the low sample number (n=3). The only potential HPV-associated discriminant taxon was *Aspergillus sulphureoviridis* (7/60) being less abundant in HPV-hr than HPV-negative samples, but this was also not significant after p value adjustment (p = 0.048, p adj = 0.765, MaAsLin2, **Supp text S4**). No taxa showed significant prevalence differences by HPV type status (p > **Supp text S7**).

### Fungal diversity analyses according to bacterial Community State Types (CSTs)

Community State Types (CSTs) were determined previously for these samples from 16S sequencing [19] and were used as variables to associate with fungal diversity in this study. Neither beta using PERMANOVA or PERMDISP nor alpha diversity were found to be significant (**Supp text S2**). However, visual inspection of the beta diversity dispersion plots suggested that CST III exhibited markedly greater variance in distance to its centroid compared to CST I and IV, supporting the use of Levene’s test to formally evaluate this observation. Levene’s test applied to centroid distances revealed significant differences in variance across CSTs (p = 0.045; Levene, **Fig. 3E**), although pairwise comparisons between CST I and IV were not significant (p adj > 0.050; pairwise Levene, **Fig. 3E, Supp text S2**). The discrepancy between the results these two variances testing methods, (Levene’s test and PERMDISP) comes from the different aspects of dispersion each method evaluates. While PERMDISP compares the mean distances to the group centroid across groups, Levene’s test compares the actual variance of those distances within each group. As such, Levene’s test provides a complementary assessment of dispersion, capable of detecting group-level heterogeneity that may not be captured by the multivariate framework used in PERMDISP.

**Figure 3.**
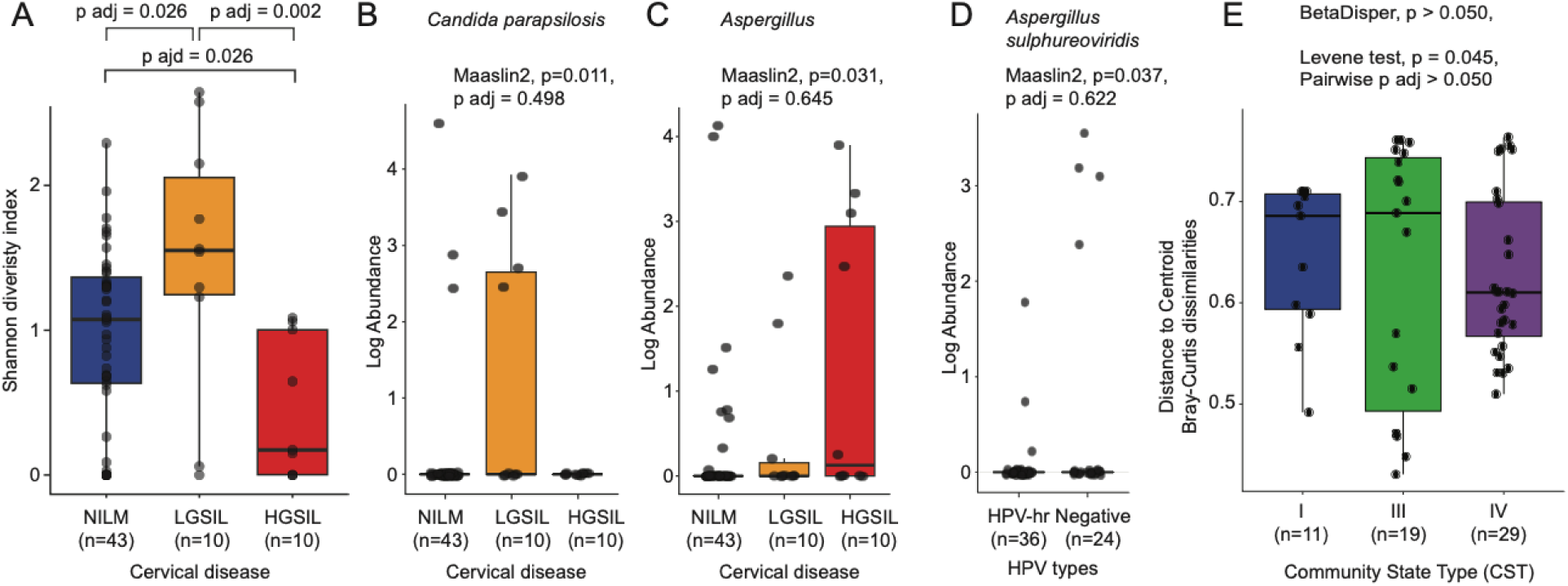
Vaginal mycobiome variation by cervical disease, HPV type, and Community State Type (CST) among Non-Pregnant population. Shannon diversity index according to cervical disease status (NILM, LGSIL, HGSIL) (**A)**. Pairwise comparisons between adjusted group means were conducted based on estimated marginal means derived from the linear model. **(B,C)** Significant taxa identified as discriminant by MaAsLin2 previous p value adjustment, across cervical disease groups, *Candida parapsilosis* **(B)**, genus *Aspergillus* **(C)**. Or across HPV types *Aspergillus sulphureoviridis(* **D)**. Distance to centroid using Bray-Curtis dissimilarity, analyzed using PERMDISP test and its pairwise form, Levene and pairwise Levene test between CSTs (**E)**.

CST groups did not show any discriminant taxa; however, the genus high the highest relative abundance, *Aspergillus*, was found to be at a lower relative abundance in CST-I compared to CST-II and IV (p = 0.133, **Supp text S4**).

In terms of taxa prevalence, no differences among CST groups were detected after p value adjustment for multiple comparisons. The species named "Fungi_sp" was found to be most prevalent in CST-IV (20.7%), absent in II and III (p = 0.039, p adj = 0.914, Fisher’s Exact Test, **Supp text S8**). This taxon is composed of 9 SHs. The most recent UNITE classification (https://unite.ut.ee/sh/) classifies the two most abundant of these SH (>23% of abundance) as an uncultured *Pyrenochaetopsis* (https://unite.ut.ee/sh/SH0994492.09FU) and *Resinicium* (https://unite.ut.ee/sh/SH1252023.09FU), among others (**Supp text S1**).

## Discussion

To our knowledge, this is the first study in the Caribbean to comprehensively characterize the cervicovaginal mycobiome in a Hispanic population across reproductive stages and cervical disease states. Using high-resolution ITS1 sequencing, we delineate the diversity, composition, and clinical associations of vaginal fungal communities, and explore their interactions with bacterial community state types (CSTs), HPV infection, and cervical lesions.

The data presented here revealed a considerable fungal diversity in the cervicovaginal environment, with about 173 species hypotheses (SH) which correspond to species level OTUs. The majority of these taxa (69%) were unique to individual samples, indicating that vaginal mycobiota exhibit substantial inter-individual variability. This pattern aligns with previous reports describing the vaginal mycobiome as a heterogeneous and individualized ecosystem [25]. Despite this diversity, only a handful of species, such as *Candida albicans, Agaricomycetes sp., Scopuloides dimorpha, Trametes cubensis* and *Hortaea werneckii* were consistently detected across multiple individuals, underscoring their ecological dominance or resilience in the vaginal niche. *H. werneckii.* is, in fact, an halophilic black yeast and the agent of tinea nigra dark macules in the palms and soles, commonly confused with skin melanoma [33]. Notably, *C. albicans*, a known opportunistic pathogen, was present in nearly half of the samples and exhibited the greatest relative abundance. Its prevalence reinforces its potential dual role as a commensal organism and a pathogenic driver under conditions of dysbiosis or immune imbalance.

The detection of non–human–associate fungi, which are very common in the outdoor air in Puerto Rico [34], including members of the Agaricomycetes class, also suggests possible environmental contributions to the vaginal mycobiota —a phenomenon increasingly reported by others [10]. Infants born in the aftermath of Hurricane Maria harbor more hurricane and asthma-related taxa, specifically *Alternaria* sp, in the nasal mycobiome [10]. This reflects the continuous influx of environmental fungi into mucosal ecosystems. The adhesion of environmental fungi to epithelial surfaces (a critical step for colonization) potentially reduces fungal and bacterial biofilm formation and alters host-microbe interactions. Such mechanisms could also operate within the vaginal environment, where airborne or dermal fungi entering the mucosa may compete with established communities for adhesion sites and nutrients, thereby influencing colonization dynamics and community stability. Moreover, environmental fungi capable of forming hyphae may enhance surface hydrophobicity and secrete hydrolytic enzymes and toxins, thereby altering local immunity and promoting inflammatory responses. These processes could indirectly impact bacterial composition, potentially driving shifts toward dysbiosis.

The presence of abundant environmental fungi in the vaginal tract, therefore, should not be dismissed as mere contamination but rather as a potential ecological force shaping the mucosal mycobiome. Our analysis revealed, however, no significant differences in alpha or beta diversity across CSTs, yet CST III displayed markedly greater fungal community variance among participants, suggesting that most detected fungi might be transient or opportunistic rather than members of a stable core mycobiome among the population. *Lactobacillus iners* the dominant species of CST III, has been associated with *Candida* pathogenic hyphal transition and biofilm formation [35], which can provide antifungal resistance [36]. This variability with the bacterial communities may indicate that the microbiota influences the stability and resilience of fungal colonization. Although no discriminant taxa were significantly associated with CSTs, the genus *Aspergillus*, a common fungus in the air, showed a trend toward lower relative abundance in CST I, a protective *Lactobacillus*-dominated environment, suggesting that beneficial bacterial communities may inhibit colonization by *Aspergillus*. Additionally, the enrichment of SH “Fungi_sp” – which included genera such as *Pyrenochaetopsis* and *Resinicium* – in CST IV, a CST often associated with dysbiosis, supports the notion that bacterial imbalance may facilitate the proliferation of otherwise rare fungal taxa. *Pyrenochaetopsis* is an environmental Ascomycete occasionally isolated from superficial human infections such as nail or skin lesions [37, 38]. *Resinicium*, a wood-decaying Basidiomycete, has no record of human pathogenicity and appears at low abundance in infant gut or nasal samples as environmental spores[39]. Both genera thus represent environmental fungi sporadically detected in human datasets, not established members of the human microbiome.

Although the reproductive stage did not significantly influence fungal alpha diversity or taxonomic composition, we observed notable differences in beta dispersion, with Non-Pregnant individuals exhibiting greater inter-individual heterogeneity compared to Pregnant women. This finding suggests that hormonal or immunological shifts during pregnancy may lead to more homogeneous fungal communities, which are often associated with *Lactobacillus*-rich epithelium in pregnancy, possibly through selective pressures favoring specific taxa. The marginally higher prevalence of *Candida tropicalis* among Pregnant participants may reflect such host-driven ecological filtering.

A major finding of this study is the reduction in fungal alpha diversity associated with high-grade squamous intraepithelial lesions (HGSIL), suggesting that diminished mycobiome diversity may accompany or contribute to the progression of advanced cervical pathology. Lower diversity could reflect a less resilient microbial community more conducive to pathogen dominance and inflammatory processes that facilitate HPV persistence and neoplastic progression. Indeed, *Candida parapsilosis* was more prevalent in low-grade lesions (LGSIL), though not statistically significant after p adjustment, suggesting potential taxon-specific associations that warrant further investigation.

HPV status (HPV-hr vs Negative) did not significantly correlate with overall fungal diversity or community composition. However, detecting *Aspergillus sulphureoviridis* with higher abundance in HPV-hr samples hints at potential interactions between specific fungal taxa and viral infection dynamics. Such interactions may occur through modulation of mucosal immunity or direct effects on epithelial integrity, production of carcinogenic mycotoxins, or as previously suggested in bacterial-fungal-viral co-infection models.

Humans, as holobionts (integrated organisms composed of host and microbial partners) are continuously exposed to environmental fungi through inhalation, ingestion, and skin contact, making such encounters inevitable. Airborne spores and hyphal fragments from genera such as *Aspergillus,* as shown here, are regularly inhaled or come into contact with the mucosal surfaces of the respiratory, gastrointestinal, and urogenital tracts ([40, 41]). These mucosal sites provide a moist and nutrient-rich microenvironments where fungi can transiently or persistently colonize. While most spores introduced into the body are efficiently cleared through mucociliary and immune mechanisms, a fraction can adhere to epithelial surfaces or germinate within the mucus layer, particularly when local immunity is compromised or the epithelial barrier is disrupted [42, 43]). Disruption of the intestinal bacterial community can compromise epithelial barrier integrity and alter immune homeostasis, creating conditions that facilitate fungal translocation beyond the gut. Bacterial dysbiosis has been shown to increase mucosal permeability and reduce colonization resistance, thereby enabling opportunistic fungi such as *Candida albicans* to traverse the gastrointestinal barrier and disseminate systemically, including to the brain [43]. The vaginal mucosa in this case may offer organic substrates and cellular debris that can sustain low-level environmental fungal growth in balance with bacterial communities, enabling the coexistence of certain environmental fungi as commensals even within asymptomatic hosts. The persistence of these fungi within mucosal niches reflects a finely tuned balance between immune recognition and tolerance. The host immune system detects fungal-associated molecular patterns while limiting unnecessary inflammation, allowing commensal species such as *Candida*, *Malassezia*, and *Aspergillus* to persist in low abundance without eliciting disease [44]. However, perturbations such as antibiotic use, any behavioral variation such as douching habits, use of detergents, or chronic inflammation can disrupt bacterial communities and mucosal immune regulation, creating ecological space for environmental fungi to expand and potentially contribute to dysbiosis. Emerging research on the aerobiome–mycobiome continuum further suggests that inhaled bioaerosols can transiently influence mucosal immune responses and microbial composition [45]. Recent hypotheses suggest that inhaled “aeronutrients”, including vitamins, fatty acids, and trace minerals, and “aeromicrobes,” such as environmental bacteria, may contribute positively to human health by supporting microbial diversity and metabolic balance within the respiratory and gastrointestinal tracts [46] with clear implications to urbanized environments and health. Building on this concept, we extend this framework to include airborne fungi as additional but potentially disruptive components of the human microbiome, including the vaginal niche. Airborne fungal spores and fragments, while ubiquitous, can interact directly with mucosal microbiomes, potentially altering their structure and function. Together, these results emphasize the intricate and dynamic relationship between environmental fungi and the host mucosa, where constant exposure, immune modulation, and microbial crosstalk govern the equilibrium between tolerance and inflammation.

## Conclusion

Our findings reveal the ecological complexity of the vaginal mycobiome and its potential influence on reproductive health and cervical disease progression. The reduction in fungal diversity associated with high-grade lesions, together with variability linked to bacterial community state types, highlights the need to consider interkingdom interactions as contributors to cervical pathogenesis. Environmental fungi within this niche may compete with resident mucosal species, shaping microbial resilience and community stability. Collectively, these insights emphasize that the vaginal mycobiome is not a passive inhabitant, but an active ecological partner shaped by host physiology and environmental exposure. Future longitudinal and mechanistic studies are essential to elucidate causal pathways and determine whether targeted modulation of fungal communities could inform new preventive or therapeutic strategies in HPV-related disease.

## Supplementary text legends

Supplementary text S1.

Summary of taxa prevalence across all individuals included in the study. The table shows the number and percentage of samples in which each taxon was detected, along with corresponding prevalence and relative abundance metrics.

Supplementary text S2.

Results of β-diversity analyses across reproductive stages and among Non-Pregnant individuals stratified by cervical disease status, HPV type, and CST. Analyses include PERMANOVA (adonis) and β-dispersion tests using Bray–Curtis and Jaccard dissimilarities, as well as Levene’s tests for CST variability.

Supplementary text S3.

Linear models of α-diversity (Shannon index) comparing reproductive stages and, among Non-Pregnant individuals, cervical disease status, HPV type, and CST. All models were adjusted for age and BMI. Pairwise comparisons were performed using estimated marginal means (EMMs) with p-value adjustment by the Benjamini–Hochberg method.

Supplementary text S4.

MaAsLin2 associations between microbial relative abundances (prevalence ≥10% and abundance ≥10 reads) and clinical variables. Only the top 10 associations per variable are displayed, ranked by absolute coefficient values.

Supplementary text S5.

Differential prevalence of taxa between study groups based on Fisher’s exact test. Only taxa meeting the inclusion threshold for prevalence and significance are shown.

Supplementary text S6.

Differential prevalence of taxa by cervical disease category (NILM, LGSIL, HGSIL) using Fisher’s exact test. Significant taxa are indicated with adjusted p-values.

Supplementary text S7.

Differential prevalence of taxa by HPV high-risk status (Positive vs. Negative) using Fisher’s exact test. Significant taxa are reported with corresponding adjusted p-values.

Supplementary text S8.

Differential prevalence of taxa across community state types (CST I–IV) using Fisher’s exact test. Significant taxa and corresponding adjusted p-values are provided.

## References

1. Ravel, J., et al., Daily temporal dynamics of vaginal microbiota before, during and after episodes of bacterial vaginosis. Microbiome, 2013. 1(1): p. 1–6.

2. Ravel, J., et al., Vaginal microbiome of reproductive-age women. Proceedings of the National Academy of Sciences of the United States of America, 2011. 108 **Suppl 1**: p. 4680–4687.

3. Godoy-Vitorino, F., et al., Malassezia globosa and Malassezia restricta are associated with high-risk HPV infections in the anogenital tract. Journal of Lower Genital Tract Disease, 2019. 22: p. PMID:–30889031.

4. Godoy-Vitorino, F., et al., Cervicovaginal Fungi and Bacteria Associated With Cervical Intraepithelial Neoplasia and High-Risk Human Papillomavirus Infections in a Hispanic Population. Front Microbiol, 2018. 9: p. 2533.

5. Bradford, L.L. and J. Ravel, The vaginal mycobiome: A contemporary perspective on fungi in women’s health and diseases. Virulence, 2017. 8(3): p. 342–351.

6. Jo, J.H., E.A. Kennedy, and H.H. Kong, Topographical and physiological differences of the skin mycobiome in health and disease. Virulence, 2017. 8(3): p. 324–333.

7. Zhang, F., et al., The gut mycobiome in health, disease, and clinical applications in association with the gut bacterial microbiome assembly. Lancet Microbe, 2022. 3(12): p. e969–e983.

8. Sharma, A., et al., Associations between fungal and bacterial microbiota of airways and asthma endotypes. J Allergy Clin Immunol, 2019. 144(5): p. 1214–1227 e7.

9. Speicher, D.J. and R.K. Aziz, Profiling the Human Oral Mycobiome in Tissue and Saliva Using ITS2 DNA Metabarcoding Compared to a Fungal-Specific Database. Methods Mol Biol, 2021. 2327: p. 253–269.

10. Acosta-Pagan, K., et al., Ecological competition in the oral mycobiome of Hispanic adults living in Puerto Rico associates with periodontitis. J Oral Microbiol, 2024. 16(1): p. 2316485.

11. Wang, R., et al., Dysbiosis in the Nasal Mycobiome of Infants Born in the Aftermath of Hurricane Maria. Microorganisms, 2025. 13(8).

12. Armstrong, E., et al., Vaginal fungi are associated with treatment-induced shifts in the vaginal microbiota and with a distinct genital immune profile. Microbiol Spectr, 2024. 12(8): p. e0350123.

13. Liu, Z., et al., Vaginal mycobiome characteristics and therapeutic strategies in vulvovaginal candidiasis (VVC): differentiating pathogenic species and microecological features for stratified treatment. Clin Microbiol Rev, 2025. 38(2): p. e0028424.

14. Lehtoranta, L., et al., Characterization of vaginal fungal communities in healthy women and women with bacterial vaginosis (BV); a pilot study. Microb Pathog, 2021. 161(Pt A): p. 105055.

15. Usyk, M., et al., TRiCit: A High-Throughput Approach to Detect Trichomonas vaginalis from ITS1 Amplicon Sequencing. Int J Mol Sci, 2023. 24(14).

16. Zhang, Y., et al., Unveiling the hidden link: fungi and HPV in cervical lesions. Front Microbiol, 2024. 15: p. 1400947.

17. Conti, H.R. and S.L. Gaffen, IL-17-Mediated Immunity to the Opportunistic Fungal Pathogen Candida albicans. J Immunol, 2015. 195(3): p. 780–8.

18. Ortiz, A.P., et al., *Incidence of Cervical Cancer in Puerto Rico*, *2001-2017*. JAMA Oncol, 2021. 7(3): p. 456–458.

19. Vargas-Robles, D., et al., The cervical microbiota of Hispanics living in Puerto Rico is nonoptimal regardless of HPV status. mSystems, 2023. 8(4): p. e0035723.

20. Tosado-Rodriguez, E., et al., Inflammatory cytokines and a diverse cervicovaginal microbiota associate with cervical dysplasia in a cohort of Hispanics living in Puerto Rico. PLoS One, 2023. 18(12): p. e0284673.

21. Gilbert, J.A., et al., The Earth Microbiome Project: Meeting report of the "1 EMP meeting on sample selection and acquisition" at Argonne National Laboratory October 6 2010. Stand Genomic Sci, 2010. 3(3): p. 249–53.

22. France, M.T., et al., VALENCIA: a nearest centroid classification method for vaginal microbial communities based on composition. Microbiome, 2020. 8(1): p. 166.

23. Pangarkar, M.A., The Bethesda System for reporting cervical cytology. Cytojournal, 2022. 19: p. 28.

24. Gonzalez, A., et al., Qiita: rapid, web-enabled microbiome meta-analysis. Nature Methods, 2018. 15(10): p. 796–798.

25. Hoggard, M., et al., Characterizing the Human Mycobiota: A Comparison of Small Subunit rRNA, ITS1, ITS2, and Large Subunit rRNA Genomic Targets. Front Microbiol, 2018. 9: p. 2208.

26. Gardes, M. and T.D. Bruns, ITS primers with enhanced specificity for basidiomycetes--application to the identification of mycorrhizae and rusts. Mol Ecol, 1993. 2(2): p. 113–8.

27. Smith, D.P. and K.G. Peay, Sequence depth, not PCR replication, improves ecological inference from next generation DNA sequencing. PLoS One, 2014. 9(2): p. e90234.

28. McMurdie, P.J. and S. Holmes, *phyloseq: an R package for reproducible interactive analysis and graphics of microbiome census data*. PloS one, 2013. 8(4): p. e61217.

29. Anderson, M.J., Permutational multivariate analysis of variance (PERMANOVA). Wiley statsref: statistics reference online, 2014: p. 1–15.

30. Oksanen, J., et al., The vegan package. Community ecology package, 2007. 10(631-637): p. 719.

31. Anderson, M.J. and D.C.I. Walsh, PERMANOVA, ANOSIM, and the Mantel test in the face of heterogeneous dispersions: What null hypothesis are you testing? Ecological Monographs, 2013. 83(4): p. 557–574.

32. Mallick, H., et al., Multivariable association discovery in population-scale meta-omics studies. PLoS Comput Biol, 2021. 17(11): p. e1009442.

33. Bonifaz, A., et al., Tinea nigra by Hortaea werneckii, a report of 22 cases from Mexico. Stud Mycol, 2008. 61: p. 77–82.

34. Rivera-Mariani, F.E., et al., Comparison of Atmospheric Fungal Spore Concentrations between Two Main Cities in the Caribbean Basin. P R Health Sci J, 2020. 39(3): p. 235–242.

35. Gaziano, R., S. Sabbatini, and C. Monari, The Interplay between Candida albicans, Vaginal Mucosa, Host Immunity and Resident Microbiota in Health and Disease: An Overview and Future Perspectives. Microorganisms, 2023. 11(5).

36. Li, H., et al., Interactions between Candida albicans and the resident microbiota. Front Microbiol, 2022. 13: p. 930495.

37. Valenzuela-Lopez, N., et al., Coelomycetous fungi in the clinical setting: morphological convergence and cryptic diversity. Journal of clinical microbiology, 2017. 55(2): p. 552–567.

38. Tsang, C.-C., et al., Diversity of phenotypically non-dermatophyte, non-Aspergillus filamentous fungi causing nail infections: importance of accurate identification and antifungal susceptibility testing. Emerging Microbes & Infections, 2019. 8(1): p. 531–541.

39. Mercer, E., et al., Divergent maturational patterns of the infant bacterial and fungal gut microbiome in the first year of life are associated with inter-kingdom community dynamics and infant nutrition. Microbiome. 12 (1): 22. 2024.

40. Esposito, M.M., S. Patsakos, and L. Borruso, The Role of the Mycobiome in Women’s Health. J Fungi (Basel), 2023. 9(3).

41. Oliveira, M., et al., Clinical Manifestations of Human Exposure to Fungi. J Fungi (Basel), 2023. 9(3).

42. Strickland, A.B. and M. Shi, Mechanisms of fungal dissemination. Cell Mol Life Sci, 2021. 78(7): p. 3219–3238.

43. Parker, A., et al., Absence of Bacteria Permits Fungal Gut-To-Brain Translocation and Invasion in Germfree Mice but Ageing Alone Does Not Drive Pathobiont Expansion in Conventionally Raised Mice. Front Aging Neurosci, 2022. 14: p. 828429.

44. Garcia-Carnero, L.C., et al., Recognition of Fungal Components by the Host Immune System. Curr Protein Pept Sci, 2020. 21(3): p. 245–264.

45. Robinson, J.M. and M.F. Breed, The aerobiome-health axis: a paradigm shift in bioaerosol thinking. Trends Microbiol, 2023. 31(7): p. 661–664.

46. Fayet-Moore, F. and S.R. Robinson, A Breath of Fresh Air: Perspectives on Inhaled Nutrients and Bacteria to Improve Human Health. Adv Nutr, 2024. 15(12): p. 100333.

